# Newcastle disease virus elevates its replication by instigating oxidative stress-driven Sirtuin 7 production

**DOI:** 10.1101/2022.10.10.511685

**Authors:** Kamal Shokeen, Sachin Kumar

## Abstract

Reactive oxygen species (ROS) accumulation inside the cells instigates oxidative stress leading to the activation of stress-responsive genes. The persistence of stress halts the cells’ antiviral response, from which numerous viruses benefit. The viral strategies for promoting stressful conditions and utilizing the induced host proteins to enhance their replication remain elusive. The present work investigates the impact of oxidative stress on NDV pathogenesis. Here, we report that the progression of NDV infection depends on intracellular ROS production. Additionally, the results demonstrate the elevation of SIRT7 levels at transcription and translational levels post-NDV infection, which in turn is associated with the positive regulation of cellular protein deacetylation. A detailed mechanistic study *in vitro* and *in ovo* was also carried out utilizing SIRT7 activity modulators to decipher the underlying role in infection, either constructive or destructive. Lastly, we concluded that the elevated expression of NDV-mediated SIRT7 protein with an enhanced activity metabolizes the NAD^+^ to deacetylase the host proteins, thus contributing to high virus replication.

**Importance:** Although the instigation of oxidative stress during NDV infection has been reported several times, the cellular stress-responsive protein’s direct function in virus replication is yet to be well understood. This study highlights the plausible stress-responsive proteins involved in viral pathogenesis while exploring the detailed molecular mechanisms of crosstalk between the activated cellular protein and the progress of the NDV replication cycle. Moreover, previous studies describing how different viruses modulate cellular stress may not fully reflect the complete picture of viral strategies. Here, we demonstrate NDV-influenced active involvement of SIRT7 activity leading to the deacetylation of host proteins. It helped us better understand the virus’s strategies to generate its numerous copies while perturbing the host cell’s standard functionality and opening up new possibilities for infection interventions.

## Introduction

Cells usually perform all their normal physiological functions in a controlled environment, and any deviation from it causes cellular stress. The high impact of stress on cells’ normal functioning makes stress management a critical factor in managing internal homeostasis. Cells respond to stress by inducing several stress-mediated signaling pathways to mitigate its effect (1). Further, the stress-activated signal transduction pathways modulate stress-responsive proteins (SRPs), maintaining protein homeostasis (2). Prolonged exposure to pathogens, mainly viruses, causes endoplasmic reticulum (ER) and oxidative stress (3–9), leading to increased stress protein production in the host. Interestingly, the virus-initiated stress has been demonstrated to exacerbate its pathogenesis, resulting in stress-induced cell death (10, 11).

Cells produce free radicals or reactive oxygen species (ROS) as a by-product of energy production (12–14). Subsequently, antioxidants remove or neutralize the ROS to maintain the proper balance (15). ROS are reactive chemical molecules that carry electron-deficient oxygen in the forms of superoxides and peroxides (16). Effective cellular sectionalization for oxygen, extensive expressions of superoxide dismutases (SODs), and glutathione S-transferase (GST) keeps the superoxide level low in the cells (17, 18). Oxidative stress-driven production of several stress-responsive proteins helps in maintaining redox homeostasis. One central regulator of oxidants exposure is nuclear factor erythroid 2– related factor 2 (Nrf2), which functions as a transcription factor driving the expression of antioxidant enzymes such as heme oxygenase-1 (HO-1) and NAD(P)H: quinone oxidoreductase 1 (NQO1) (19). In addition, heat shock proteins (majorly HSP90), which are ATP-dependent chaperones, are also reported to be actively involved in oxidative stress (20, 21).

The most common and potent stressors are viruses, which trigger various pathways to create a stressed environment (22). Studies demonstrate that viruses, including human immunodeficiency (23, 24), dengue (25, 26), influenza (27), and hepatitis (7, 28) virus, modulate oxidative stress and specific oxidant-sensitive pathways. Newcastle disease virus (NDV) causes a highly infectious disease in avian species and has also been reported to induce cellular oxidative stress (29–31). Studies showed oxidative imbalance and decreased antioxidant activity, including SODs, GST, and catalase, in chickens infected with NDV (32). Furthermore, it has also been observed that the antioxidant defense system is impaired following NDV infection, and supplementation of vitamin E helps in restoring it (30, 33, 34). Accumulating ROS and oxidative damage play a crucial role in viral diseases; thus, establishing their precise part is a prerequisite to revealing new insights into cell defense strategies while identifying new therapeutic targets.

Sirtuins (SIRTs), comprising seven different proteins, are a family of nicotinamide adenine dinucleotide (NAD)-dependent histone deacetylases identified as essential for cell survival (35). They are ubiquitously expressed and evolutionarily conserved that are actively involved in DNA repair, autophagy, aging, cardiovascular diseases, and several neurodegenerative diseases (36). SIRT1, SIRT6, and SIRT7 are primarily located in the nucleus; SIRT2 is a cytoplasmic protein, whereas SIRT3, SIRT4, and SIRT5 are distributed in mitochondria (35, 37–40). Additionally, they are linked with oxidative stress-driven cellular responses triggered to curtail the stress via deacetylation of various substrates (41–43). SIRTs are reported to be involved in virus infection; several reports pointed them as essential defense elements against several viruses (44–46), while others suggested their constructive participation in viral pathogenesis (47–50).

SIRT7 is the least explored protein among all members as its enzymatic activity, physiological function, and target proteins remain elusive. It belongs to class III histone deacetylase (HDAC) and is a critical regulator of host gene expression (51). However, recent studies have revealed its deacetylase function on several non-histone proteins, including ATM (52), p53 (53), GATA4 (54), and PAF53 (55). Presumably, it may restrict as well as enhance the virus replication via deacetylation of a different set of proteins (46, 56). However, recent studies focus on deciphering its role as an essential cellular regulator and utilizing it as a therapeutic target to explore future therapies for multiple diseases (57). SIRT7-mediated deacetylation and its involvement in virus replication are yet to be understood. Most importantly, the molecular mechanisms of crosstalk between SIRT7 protein and the virus replication cycle are not well explored. Therefore, many previous studies describing how different viruses modulate the SIRT7 expression may not fully reflect the bigger picture of their participation in viral pathogenesis.

The present study primarily focuses on elucidating the role of intracellular ROS in NDV infection by studying its replication kinetics in normal and oxidative-stressed conditions. The study aims to point out the part of stress-responsive proteins in viral pathogenesis while exploring the underlying mechanism behind.

## Materials and Methods

### Cell culture and Agents

Chicken embryo fibroblast cells (DF-1) (UMNSAH/DF-1, ATCC® CRL-12203™) were cultured in Dulbecco’s modified eagle’s media (DMEM), containing 10% fetal bovine serum (FBS) and 1% antibiotic-antimycotic solution (HiMedia, India). The monolayer was maintained in a humidified 37°C incubator under 5% CO_2_. Dimethyl furamate (DMF), pyrrolidine dithiocarbamate (PDTC), N-Acetyl-L-cysteine (NAC), lithium chloride (LiCl), β-nicotinamide mononucleotide (β-NMN), nicotinamide (NAM), and β-nicotinamide adenine dinucleotide (β-NAD) were procured commercially (Sigma, USA, and SRL, India). The stock solutions were prepared in dimethyl sulfoxide (DMSO) or PBS and diluted in a culture medium. The cell-permeant ROS probe 2’-7’-Dichlorodihydrofluorescein diacetate (DCFH-DA) (Cat# D6883, Sigma, USA) was also diluted in DMSO.

### Plasmid Transfection

To achieve the heterologous expression of SIRT7, a gene SIRT7 (C-terminally 3xFLAG tagged) under the control of a CMV promoter in pcDNA3.1 was commercially synthesized (GenScript, USA). For transfection experiments, DF-1 cells were seeded on 6-well plates and transfected with two μg of expression vector using Lipofectamine 2000 (Invitrogen, USA) according to the manufacturer’s instructions.

### Virus Infection and quantification

The mesogenic strain NDV R2B was used for conducting all the experiments, while the recombinant NDV expressing available in the laboratory was utilized for the fluorescent-based study (3, 58). The amplification, quantification, and viral kinetic assays were performed in DF-1 cells. Two different conditions with DMF (12 hrs pre-treatment) and PDTC (12 hrs post-treatment) were used to examine the effects of oxidative stress on NDV infectivity. For pre-treatment experiments, cells were seeded with the density of 5×10^5^ cells/well in 6-well plates. After six hours, the cells were pre-treated with DMF (1 μM) and incubated for another 12 hrs in a humidified 37°C incubator under 5% CO_2_. After incubation, the monolayer was infected with 0.01 MOI of NDV; subsequently, cells and supernatant were collected at 24, 48, and 72 hrs post-infection. For post-treatment experiments, the cell monolayer was infected with 0.01 MOI of NDV, and the cells were treated with PDTC (5 μM) 12 hrs post-infection. Finally, cells and supernatant were collected at 24, 48, and 72 hrs post-infection. The viral quantification was done using TCID_50_ and plaque assay as per the standard protocol (59–61). Similarly, for the SIRT7 activity assessment study, cells were subjected to two conditions with β-NMN (1mM) (12 hrs pre-treatment) for activation and NAM (1mM) (12 hrs post-treatment) for inhibition. Additionally, for *in ovo* study, nine-day-old specific-pathogen-free embryonated eggs were used for virus inoculation and compound treatments. Allantoic fluid and embryo tissue samples were collected 48 hrs post-NDV inoculation.

### Measurement of ROS levels

The generation of intracellular ROS was determined using DCFH-DA dye (Sigma, USA). Cells were seeded in 96-well plates and incubated overnight in a humidified 37°C incubator under 5% CO_2_. DMF pre-treatment and PDTC post-treatment assays, along with virus infection, were done, as discussed earlier. At 24, 48, and 72 hrs post-NDV infection, the growth media was aspirated from each well, and cells were incubated with DCFH-DA dye at a final concentration of 10 μM for 30 min at 37°C in the dark. Post-incubation, cells were washed three times with PBS, and subsequently, the fluorescence intensity was measured using the Tecan infinite 200Pro device with excitation at 493 nm and emission at 523 nm. Mean fluorescence intensity was calculated to plot the intensity graph. Changes in the emitted fluorescence intensity post-virus infection and compound treatment allowed interpretations for the production of ROS in mitochondria.

### Gene Expression Study

Total RNA was extracted from the cells reserved from compound treatment and virus infection experiments using RNAiso Plus reagent (TaKaRa, Japan) to analyze gene expression by quantitative PCR. 1000 ng of isolated RNA were subjected to cDNA synthesis using the high-capacity cDNA reverse transcription kit (Thermo Scientific, USA), followed by PowerUp SYBR Green Master Mix (Applied Biosystems, USA) according to the manufacturer’s recommended cycling conditions. Transcript levels were quantified using host and NDV gene-specific primers and normalized to GAPDH levels. With final normalization to untreated and uninfected controls, relative fold change values were calculated by the comparative cycle threshold (2^−ΔΔCT^) method (62).

### Immunoblot Analysis

DF-1 cell lysates were collected in RIPA buffer, and an equal amount of protein per sample was subjected to SDS-polyacrylamide gel electrophoresis. The proteins were transferred to a 0.4 μm nitrocellulose membrane and subsequently blocked in 5% skimmed milk containing 0.1% Tween-20 in TBS for 2 hrs at room temperature. The membrane was then probed with target protein-specific primary antibodies overnight at 4°C, followed by horseradish peroxidase (HRP)-conjugated secondary antibodies (Cell Signaling Technology, USA) for one hr at room temperature. Lastly, the blots were detected using ECL Detection Reagent (BioRad, USA), and signals were detected by chemiluminescence. Primary antibodies against chicken NDV proteins (available in the laboratory (3, 63, 64)), β-actin (Invitrogen, USA), sirtuin 7, and acetylated lysine (Cell Signaling Technology, USA) were used.

### Flow cytometric Analysis

DF-1 cells were cultured in a 6-well plate at 5×105 cells/well for flow cytometric assays and subjected to virus infection and compound treatment. At the end of the infection and treatment, cells were harvested, and the cell pellet was washed with ice-cold PBS twice for the complete removal of media. Finally, cells were resuspended in 2% FBS-PBS and analyzed by a CytoFLEX flow cytometer (Beckman Coulter, USA). The final results from three independent experiments were averaged and shown in the histogram.

### NAD^+^ and NAM detection

Nine-day-old chicken embryonated embryos were inoculated with 100 μl of NDV with 10^6^ pfu/ml and subsequently incubated in a 37°C incubator for two days. Equal weights (500 mg) of embryonic tissue samples were collected from control and infected groups. After the trituration in an equal volume of plain DMEM, the samples were vortexed thoroughly to mix the cells evenly. Tissue samples were added with pre-chilled 10% perchloric acid (HClO_4_) at a 1:10–50 ratio (tissue weight: HClO_4_ volume) and kept on ice for 15 min. After centrifuging the samples at 22,000g at 4°C for 10 min, the supernatant was transferred to a new tube and subsequently added with 3M potassium carbonate (K_2_CO_3_) and vortex rigorously. The mixture was then incubated on ice for 15 minutes and centrifuged at 22,000xg at 4°C for 10 minutes. Finally, the supernatant was transferred to HPLC glass vials. Tissue NAD^+^ and NAM levels were determined using a High-Performance Liquid Chromatography (HPLC) system (Shimadzu, Kyoto, Japan) with a C18 column (5 μm, 4.6×250 mm) by absorbance at 261 nm. Mobile phase A consisted of 0.05 M phosphate buffer (pH 7.0), and mobile phase B was 100% methanol. The following linear gradient was run for 30 min at a flow rate of 1 μl/min with the running program already described previously (65). Multiple known concentrations of NAD^+^ and NAM were prepared in PBS and used to plot the standard curve. Lastly, the injection volume was 100 μl per sample, and all the samples were kept at 4°C in the autosampler tray prior to injection.

### Statistical Analysis

Results were expressed as mean ± standard deviation (SD). The data obtained were statistically analyzed using a two-way analysis of variance (ANOVA) in GraphPad Prism software. The statistical significance among different groups was indicated as *, **, and *** where p < 0.05, p < 0.01, p < 0.001, respectively.

## Results

### Intracellular ROS production post-NDV infection

The cytotoxicity assessment of DMF (ROS inducer) and PDTC (ROS scavenger) in DF-1 cells was performed at four different concentrations (1, 5, 10, and 20 μM) using 3-(4,5-dimethylthiazol-2-yl)-2,5-diphenyltetrazolium bromide (MTT) calorimetry assay. There were no impacts on the growth of the DF-1 cell line till five μM, whereas the maximum toxicity recorded at 20 μM concentration was about 20% inhibition value for DMF and 50% for PDTC (Figures 1A and 1D). Therefore, one μM of DMF and five μM of PDTC were chosen as the optimal concentration for intracellular ROS induction control. Subsequently, intracellular ROS induced upon NDV infection were detected using fluorescent DCFH-DA dye. A persistent increase in fluorescence intensity was observed, up to 150%, till 72 hrs post-NDV infection (Figure 1B). This effect was almost equivalent to DMF-treated positive control cells. Furthermore, LiCl, a previously described antiviral drug for NDV, was used to observe the reversion of ROS levels inside the cells. The increased fluorescence intensity was diminished to 65% on inhibiting NDV replication via LiCl treatment (Figure 1C). In contrast, known ROS scavengers PDTC and NAC were also used to confirm the modulation in NDV-infected cells. The post-treatment of NDV-infected cells with these drugs exhibited an impairing effect on the NDV-induced ROS, reaching more than 50% in PDTC and almost 100% by NAC, respectively (Figures 1E and 1F).

**Figure 1.**
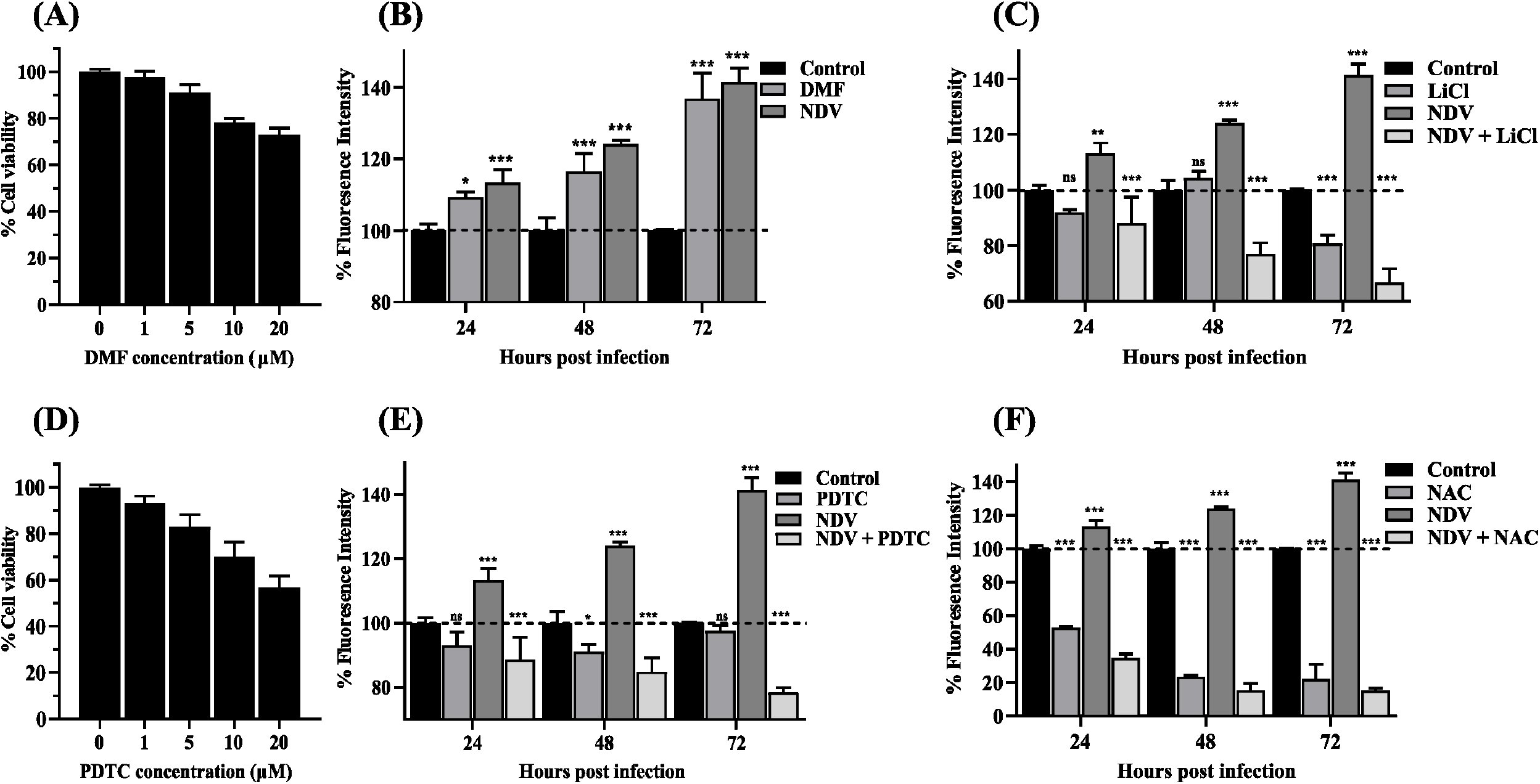
Intracellular ROS production post-NDV infection. Cytotoxicity of drugs in DF-1 cells using MTT assay. Graph representing the percentage of cell viability after 24 hrs treatment with increasing concentrations of DMF (0-20 μM) **(A)** and PDTC (0-20 μM) **(D).** Generation of intracellular ROS due to NDV with DMF **(B)**, with LiCl **(C)**, with PDTC **(E)**, and with NAD **(F)** through fluoroscopic measurements determined using DCFH-DA dye. Values represent the mean fold change ± S.D. Statistical analyses were performed using a two-way ANOVA (p<0.05, p<0.001, and p<0.001 are described as *, **, and ***).

### Replication kinetics of NDV in oxidative condition

Cells were pre-incubated with DMF for 12 hrs mimicking the oxidative stressed environment to examine the possible participation of ROS generation in NDV infection. The gene and protein expression experiments showed that DMF-treated cells had higher virus transcript and protein levels than untreated groups (Figures 2A and 2B), and the viral kinetic profile was consistent at all time intervals. The overall virus titer was also determined and compared with untreated groups using TCID_50_ and plaque assay with similar conditions. The maximal response was noted at 72 hrs post-NDV infection, where the mean viral load was 8.33 log_10_ TCID_50_ per ml in stressed cells, compared to 6.83 log_10_ TCID_50_ per ml in normal cells (Figure 2C). Similarly, more than 100% of plaques forming units (PFU) than control groups were observed at 48 and 72 hrs post-NDV infection (Figure 2D).

**Figure 2.**
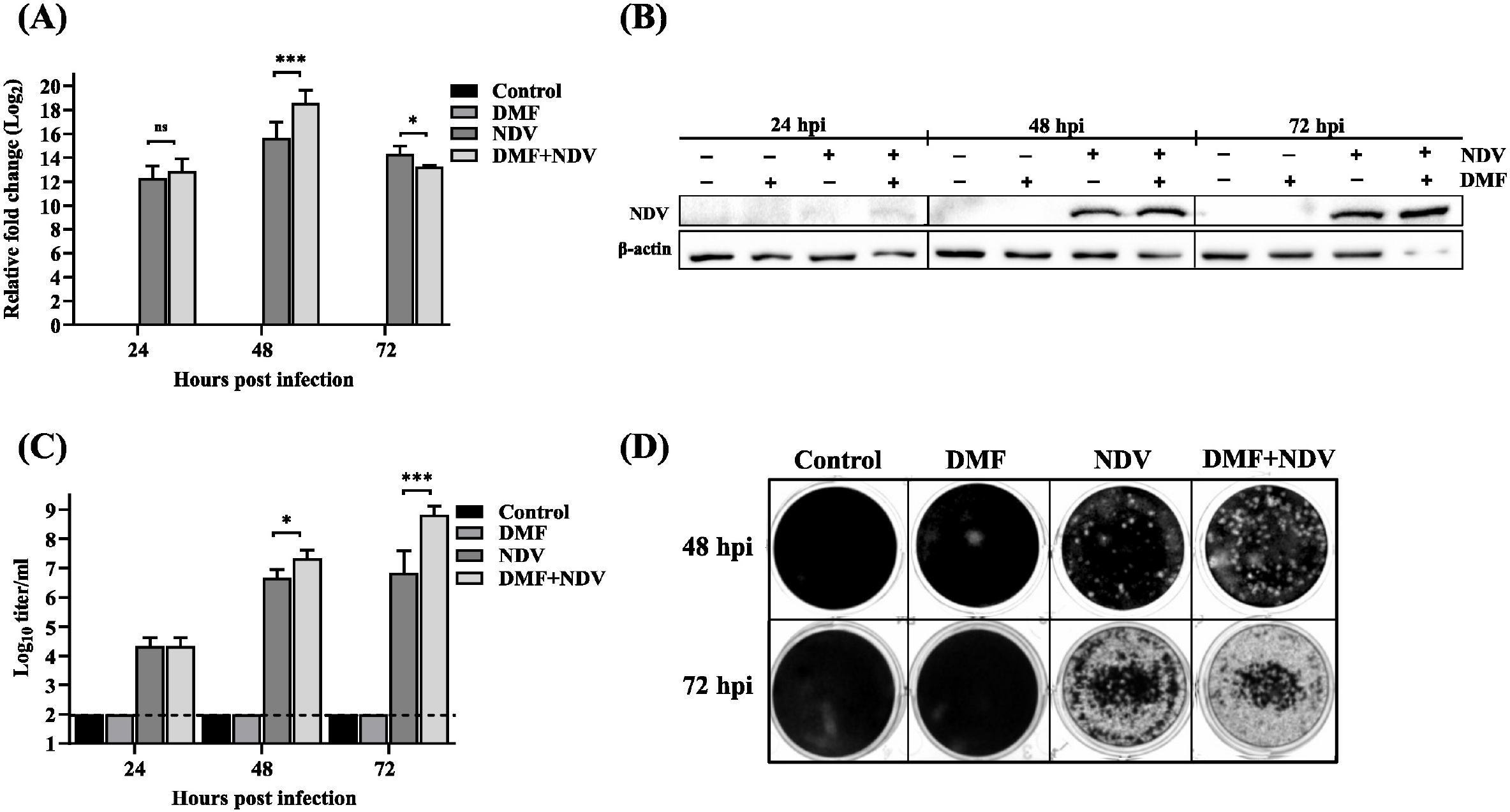
Replication kinetics of NDV in oxidative condition. The graph shows the relative fold change in NDV N gene mRNA in DF-1 cells with and without DMF treatment **(A)**, with GAPDH as an internal control for normalization. Immunoblot analysis of NDV and β-actin in DF-1 cells when treated with DMF (1 μM) at different time intervals **(B).** Assessment of NDV titer after treatment with DMF using TCID_50_ assay **(C)**. Assessment of NDV titer after treatment with DMF using plaque assay **(D).** Values represent the mean fold change ± S.D. Statistical analyses were performed using a two-way ANOVA (p<0.05, p<0.001, and p<0.001 are described as *, **, and ***).

Contrarily, the effects of reduction of ROS generation on NDV pathogenesis were also investigated, where cells were post-treated with ROS scavenger drug. Post-NDV infection PDTC treatment significantly inhibited virus replication (up to 8-fold) and protein production (Figures 3A and 3B). In congruence with the above results, the TCID_50_ and plaque assay subsequently revealed the reduced titer of NDV in PDTC-treated cells. The mean viral load was reduced to 5.6 log_10_ TCID_50_ per ml from 6.8 log_10_ TCID_50_ per ml in control groups at 48 hrs post-infection. However, at 72 hrs, the reduction in NDV titer did not reach the statistical significance level (Figure 3C). Similarly, a notable decline of NDV plaques to almost 100% and around 50% was observed compared to control at 48 and 72 hrs, respectively (Figure 3D).

**Figure 3.**
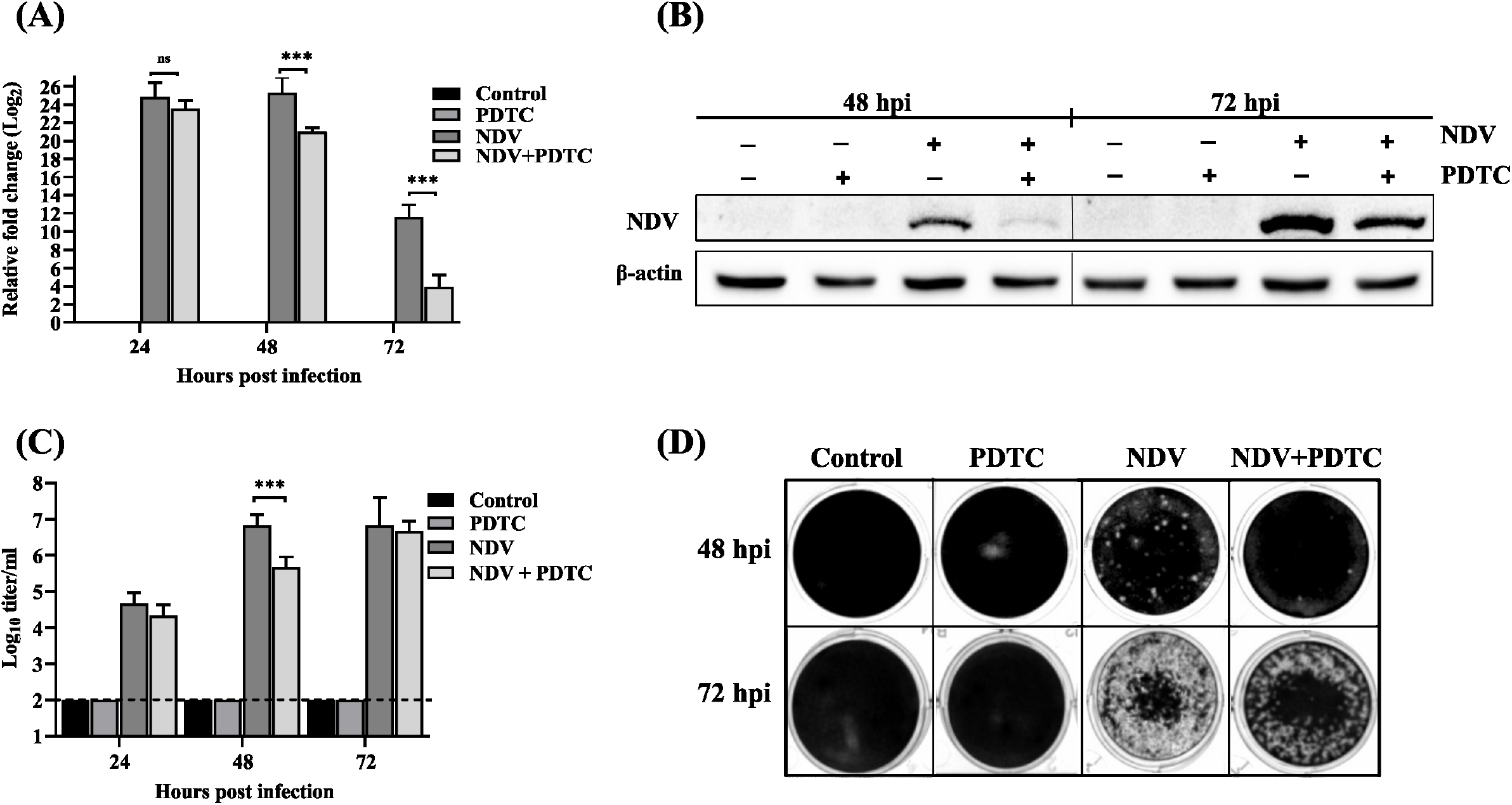
NDV infection after inhibiting intracellular ROS production. The graph shows the relative fold change in NDV N gene mRNA in DF-1 cells with and without PDTC (A) treatment, with GAPDH as an internal control for normalization. Immunoblot analysis of NDV and β-actin in DF-1 cells when treated with PDTC (1 μM) at different time intervals **(B).** Assessment of NDV titer after treatment with PDTC using TCID_50_ assay **(C)**. Assessment of NDV titer after treatment with PDTC using plaque assay **(D).** Values represent the mean fold change ± S.D. Statistical analyses were performed using a two-way ANOVA (p<0.05, p<0.001, and p<0.001 are described as *, **, and ***).

### Modulation of oxidative stress-response genes post-NDV infection

The expression patterns of crucial oxidative stress-responsive genes in NDV-infected cells were determined and compared to mock-infected DF-1 cells. At first, the relative fold change of the NDV N gene was calculated at 24, 48, and 72 hrs to confirm the progress of virus replication (Figure 4A). Subsequently, expression profiles of Nrf2, HO-1, SOD-1, and Hsp90b were studied with or without NDV at different time points. The Nrf2 mRNA expression was decreased from 1.0 to 0.3-fold (Figure 4B), followed by HO-1 inhibition from 1.0 to 0.25-fold after 72 hrs of NDV infection (Figure 4C). At the same time, SOD-1 was upregulated up to three-fold at 24 hrs post-infection but showed a marked drop at 48 hrs when NDV replication reached its peak (Figure 4D). Concurrently, Hsp90b over-expression was progressive and time-dependent, getting a 4-fold increment (Figure 4E). Furthermore, mRNA variation levels of sirtuin family genes (SIRT1, SIRT3, SIRT5, and SIRT7) instigated by NDV were also calculated. NDV infected cells demonstrated significant up-regulation of SIRT1 (from 1.0 to 4.5-fold at 24 hrs) and SIRT7 (from 1.0 to 11.0-fold at 48 hrs) expression levels as well as substantial down-regulation of SIRT3 (from 1.0 to 0.22-fold at 48 hrs) and SIRT5 (from 1.0 to 0.15-fold at 72 hrs) expression levels compared to the control groups as measured by qPCR (Figures 4F-I).

**Figure 4.**
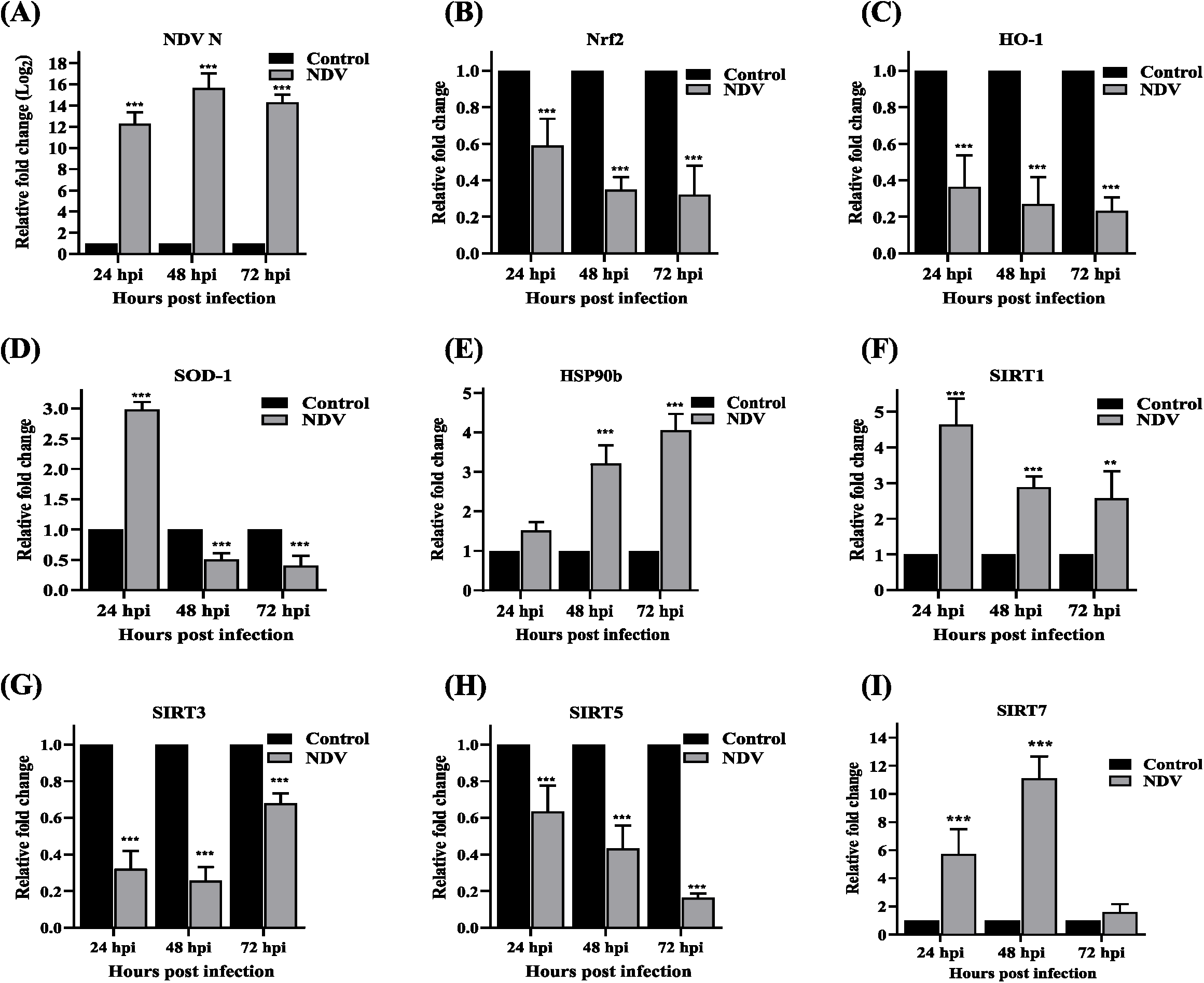
Modulation of oxidative stress-response genes post-NDV infection. The graphs show the relative fold change in the NDV N **(A)**, Nrf2 **(B)**, HO-1 **(C)**, SOD1 **(D)**, HSP90b **(E)**, SIRT1 **(F)**, SIRT3 **(G)**, SIRT5 **(H)**, and SIRT7 **(I)** transcript levels after 24, 48, and 72 hours post-infection with NDV and DMF was used as a positive control for oxidative stress. The gene-specific primers were used for the analysis, with GAPDH as an internal control for normalization. Values represent the mean fold change ± S.D. Statistical analyses were performed using a two-way ANOVA (p<0.05, p<0.001, and p<0.001 are described as *, **, and ***).

### NDV modulated SIRT7 production

The SIRT7 protein modulation upon NDV infection was studied in a time-dependent manner. The results showed dramatic overexpression of SIRT7 upon NDV infection at 48 hrs (Figure 5A). Additionally, cells were treated with varying concentrations of DMF (1 and 5 μM) for 24 hrs and subsequently subjected to western blot analyses, which showed that high levels of SIRT7 were induced in oxidatively stressed cells (Figure 5B). In contrast, the effects of PDTC on SIRT7 mRNA turned out to be around 80% lower (from 1.0 to 0.22-fold at 24 hrs) in treated cells at 24hrs (Figure 5C). To understand the effect of PDTC treatment on NDV-induced SIRT7, the expression of SIRT7 protein was studied in the presence of NDV and in combination with PDTC. While the up-regulation of SIRT7 at the transcriptional and translational level was observed in NDV alone, the PDTC post-treatment data showed that SIRT7 expression induced by NDV was substantially suppressed (Figures 5D and 5E). For the heterologous production, pcDNA3.1 expressing SIRT7-cFLAG was utilized. The plasmid was verified using restriction digestion analyses (Figure 5F); meanwhile, the expression was confirmed by western blot, which showed the presence of SIRT7 protein when detected with an anti-FLAG antibody (Figure 5G). Next, to examine the supportive effects of SIRT7 on NDV replication, DF-1 cells were transfected with the plasmid 12 hrs before NDV infection. The immunoblot results showed that the NDV protein production was considerably higher in SIRT7-overexpressing cells from empty vector-transfected cells (Figure 5H).

**Figure 5.**
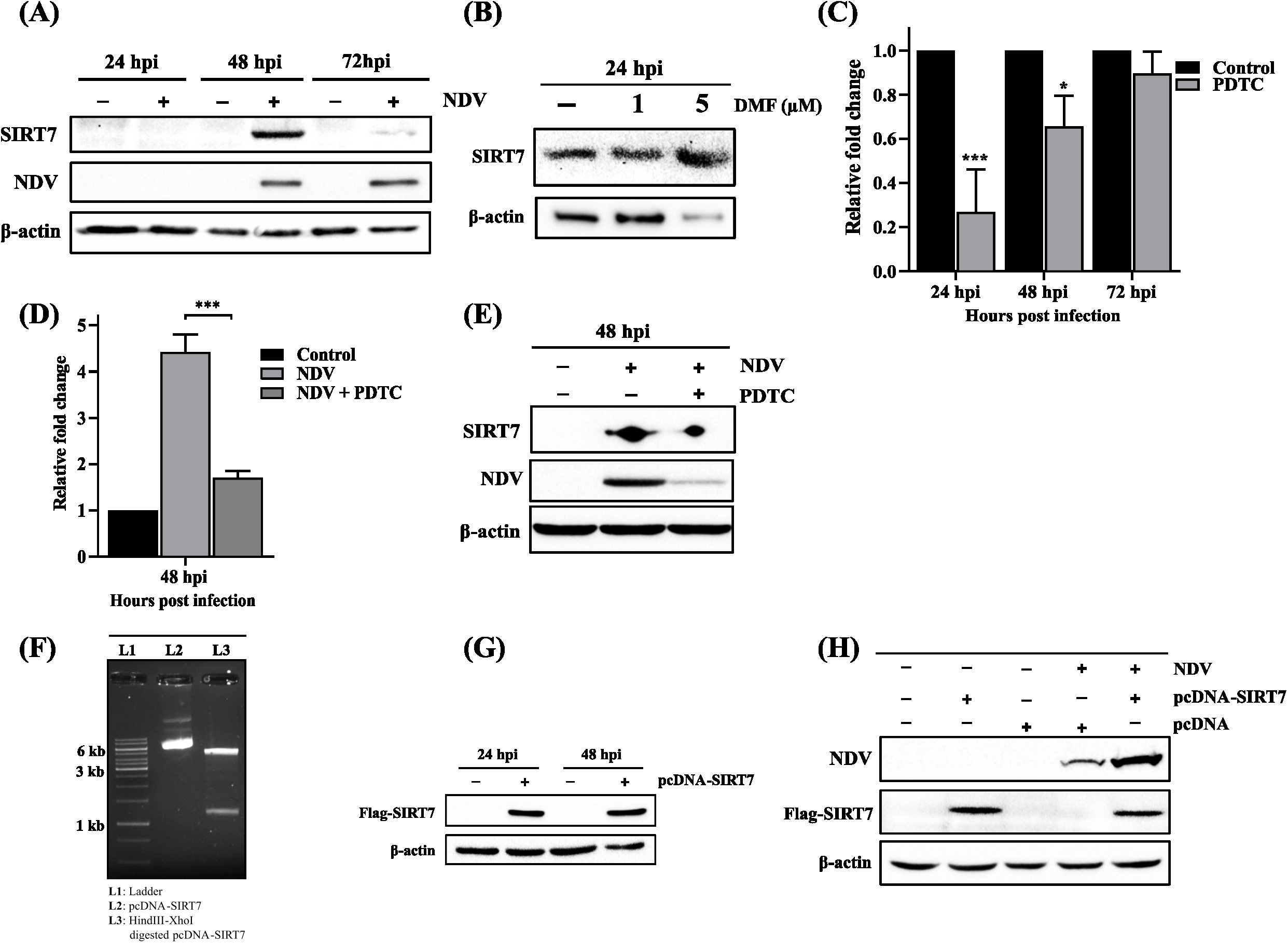
Modulation of SIRT7 production mediated by oxidative stress post-NDV infection. Immunoblot analysis of SIRT7, NDV, and β-actin in DF-1 cells after 24, 48, and 72 hours post-infection with NDV **(A)**. Immunoblot analysis of SIRT7 and β-actin after treatment with DMF (1 and 5 μM) for 24 hrs **(B)**. The graph shows the relative fold change in SIRT7 gene mRNA influenced by PDTC (5 μM) for different time intervals **(C)**. The graph shows the relative fold change in SIRT7 gene mRNA in DF-1 cells with and without the treatment of PDTC **(D)**. GAPDH was used as an internal control for normalization. Immunoblot analysis of SIRT7, NDV, and β-actin in DF-1 cells with and without the treatment of PDTC **(E)**. Agarose gel image showing digestion of the pcDNA-SIRT7 plasmid with HindIII and XhoI restriction enzymes **(F)**. Immunoblot image showing the expression of SIRT7 protein, detected by FLAG antibody, after transfection with a pcDNA-SIRT7 plasmid **(G)**. Immunoblot shows the production of NDV protein along with SIRT7 and β-actin after over-expressing SIRT7 protein by transfection **(H)**. Values represent the mean fold change ± S.D. Statistical analyses were performed using a two-way ANOVA (p<0.05, p<0.001, and p<0.001 are described as *, **, and ***).

### SIRT7 activity involvement in NDV infection

To decipher the biological significance of SIRT7, global acetylation level was detected using an anti-acetylated lysine antibody in NDV-infected DF-1 cells. Interestingly, the virus infection led to a significant reduction of acetylation levels in a time-dependent manner (Figure 6A). Subsequently, a similar depletion of acetylated proteins was also observed when infected with different MOI of NDV (0.01, 0.1, and 1 MOI) (Figure 6C). These changes in acetylation level were confirmed by infecting the cells with heat-inactivated NDV particles. Compared with the NDV-infected cells where the acetylated protein has completely vanished, it is evident that the deacetylation effect on cellular proteins was reverted to normal conditions in the case of inactivated NDV (Figure 6D). Furthermore, the deacetylation activity of SIRT7 was tested at 24 and 48 hrs in SIRT7 overexpressed cells. From the western blot analyses, the presence of excess SIRT7 in cells showed a similar trend in acetylation level compared to NDV-infected groups (Figure 6E). Lastly, to further validate the current findings, β-NMN, a known SIRT7 activity enhancer, was utilized to monitor the acetylated protein levels. The addition of β-NMN in cells showed lower levels, which are in corroboration with previous results (Figure 6F).

**Figure 6.**
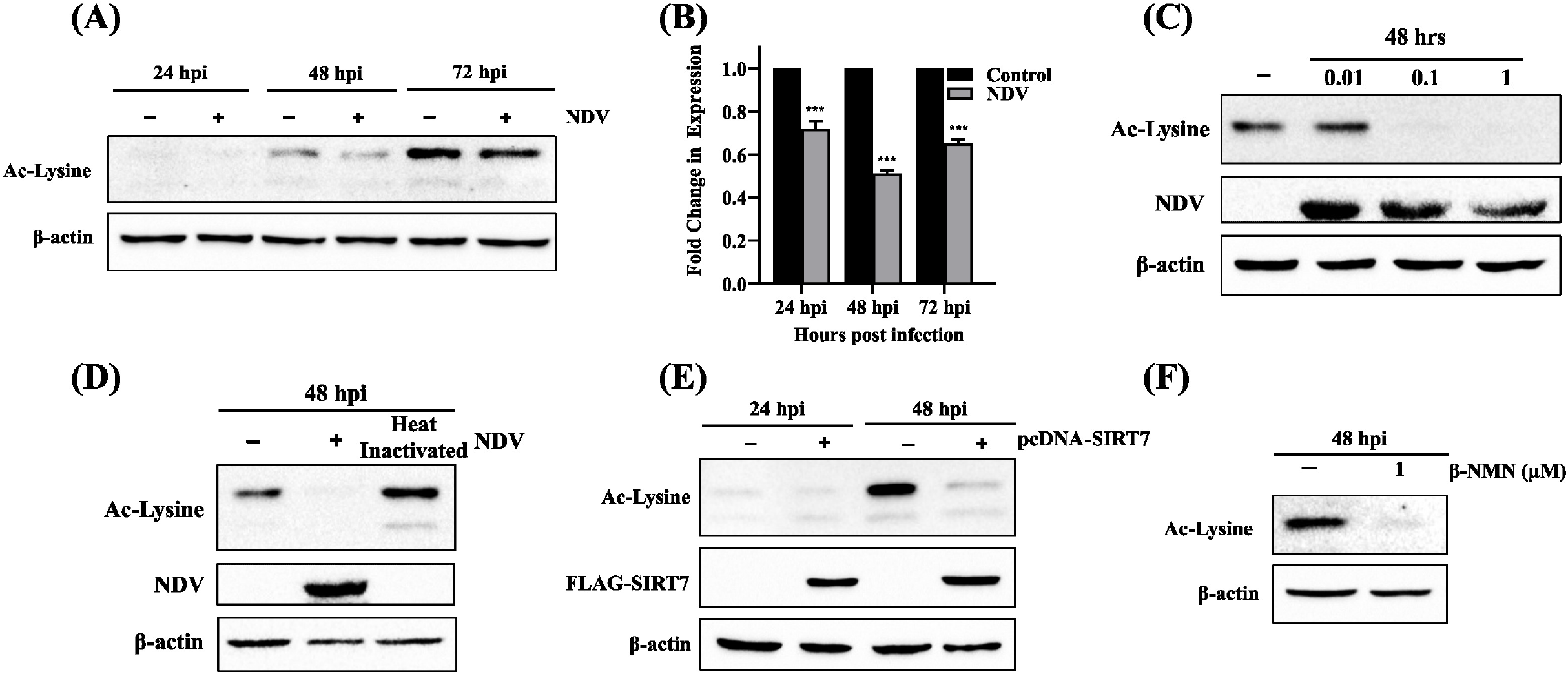
SIRT7 activity involvement in NDV infection. Immunoblot analysis of acetylated-lysine (representing acetylated-proteins) and β-actin in DF-1 cells after 24, 48, and 72 hours post-infection with NDV **(A)**. The graph shows the fold change in expression instigated by NDV in acetylated lysine **(B)**. Immunoblot image showing the effect on acetylated-lysine levels in DF-1 cells infected with multiple MOIs **(C)**. Immunoblot image comparing the acetylated-lysine levels in NDV infected cells with heat-inactivated NDV infected cells **(D)**. Immunoblot image showing the effect of SIRT7 overexpression on acetylate-lysine after transfection **(E)**. Immunoblot image showing the activity of SIRT7 on acetylated proteins after treatment with β-NMN for 48 hrs **(F)**.

To get a clear picture of the exact function SIRT7 plays in NDV replication, DF-1 cells were pre-incubated with β-NMN for 8 hrs before infection. The immunoblot images showed that β-NMN treated cells exhibit higher viral protein production (Figure 7A). On the contrary, the post-treatment of NAM, SIRT7 activity feedback inhibitor, was done 8 hrs after infection. The results demonstrated the virus inhibitory effects of NAM in NDV-infected cells, with a notable decrease in viral proteins (Figure 7B). Effects of the compounds were further validated by visualizing the number of cells expressing the NDV-encoded green fluorescent protein (GFP). As expected, cells with β-NMN treatment exhibit relatively more GFP fluorescence. In contrast, cells with NAM treatment showed reduced fluorescence intensity, representing the NDV replication proportion in respective groups (Figure 7C). To quantify the GFP expressed by NDV, infected cells were analyzed for GFP intensity by flow cytometry. Expectedly, the GFP positive cells were more frequent in β-NMN treated infected cells (49.78%) but were rare in NAM treated cells (23.99%) than NDV control (41.50%), confirming the participation of these compounds in viral pathogenesis quantitatively (Figure 7D).

**Figure 7.**
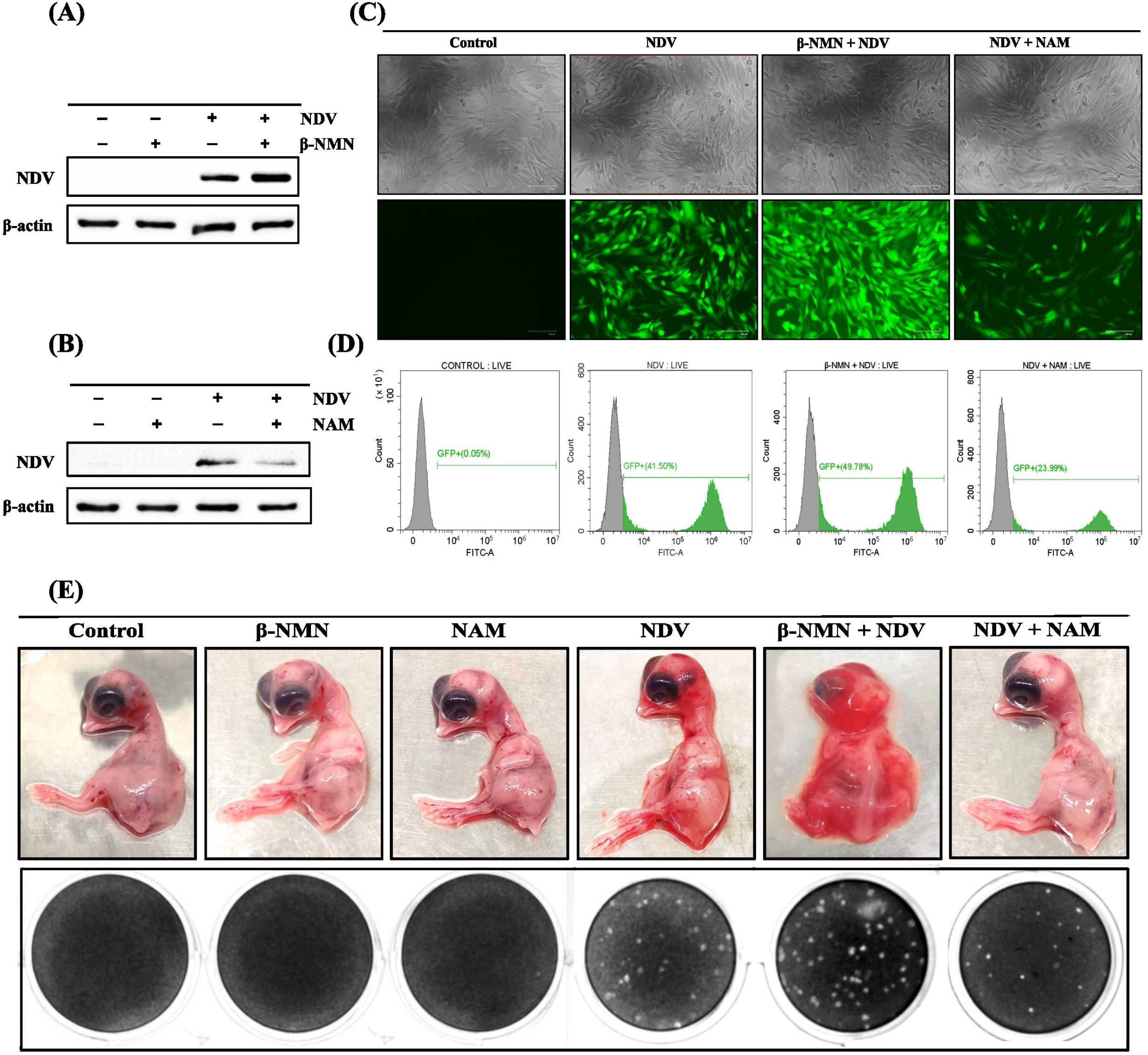
*In vitro* and *in ovo* SIRT7 activity assessment. Immunoblot analysis of NDV and β-actin in DF-1 cells with and without β-NMN treatment **(A)**. Immunoblot analysis of NDV and β-actin in DF-1 cells with and without NAM treatment **(B)**. Impact of β-NMN and NAM treatment on GFP expressing recombinant NDV using microscopy imaging and flow cytometric analysis **(C)(D)**. Gross lesions and hemorrhages in 9-days-old chicken embryos after 48 hrs of NDV infection in the absence and presence of β-NMN and NAM, respectively. Quantification image of NDV titer in allantoic fluid from embryonated eggs using plaque assay **(E).**

Given a strong correlation between the NDV infection and SIRT7 activity, similar treatment studies were also performed *in ovo* system where nine-day-old embryonated eggs were pre-and post-treated with the respective compounds. NDV-infected chicken embryos without treatment were characterized with hemorrhages, which were comparatively more in β-NMN pre-treated embryos. The embryos with β-NMN pre-treatment and NDV inoculation died in 48 hrs and were found completely liquified, as shown in Figure 7E; however, the embryos with compound treatment alone showed no lesions. NAM post-treated NDV-infected embryos looked very similar to their respective control groups. The virus titer in the allantoic fluid collected from the embryos was also determined by standard plaque assay. The NDV titer in the β-NMN treated embryos was calculated to be 3.95×10^8^ pfu/ml, significantly higher than virus control (1.95×10^8^ pfu/ml). The NDV titer in NAM post-treated embryos was calculated as 0.9×10^8^ pfu/ml (Figure 7E).

### NAD^+^ and NAM estimation in NDV-infected embryos

Considering the involvement of NAD^+^-dependent SIRT7 in NDV infection, the NAD^+^ levels in NDV-infected DF-1 cells were compared with control cells. Firstly, the known concentrations of NAD^+^ and NAM were injected into the HPLC column to determine the peak retention time; the chromatograms are shown in Figure S1. The NAD^+^ and NAM peaks were obtained around 16.5 and 18.4 min. Next, the DF-1 cell samples were distributed in two halves, with only one half supplemented with NAD^+^ to point the specific NAD^+^ peak out of multiple peaks; the respective chromatograms are shown in Figure S2. The DF-1 cells data showed a gradual decrease in the NAD+ concentrations in infected cells, with a minimal difference of around 11% at early time intervals and reaching a 60% reduction post 72 hrs of infection (Figure 8A). The chromatograms of uninfected and NDV-infected cells are shown in Figure S3. Afterward, the results were validated by estimating the NAD^+^ concentration along with NAM in NDV-infected embryo tissues. *In ovo* study results showed 20% lower NAD+ levels in the tissues, whereas NAM levels were estimated to be 130% compared to control embryos.

**Figure 8.**
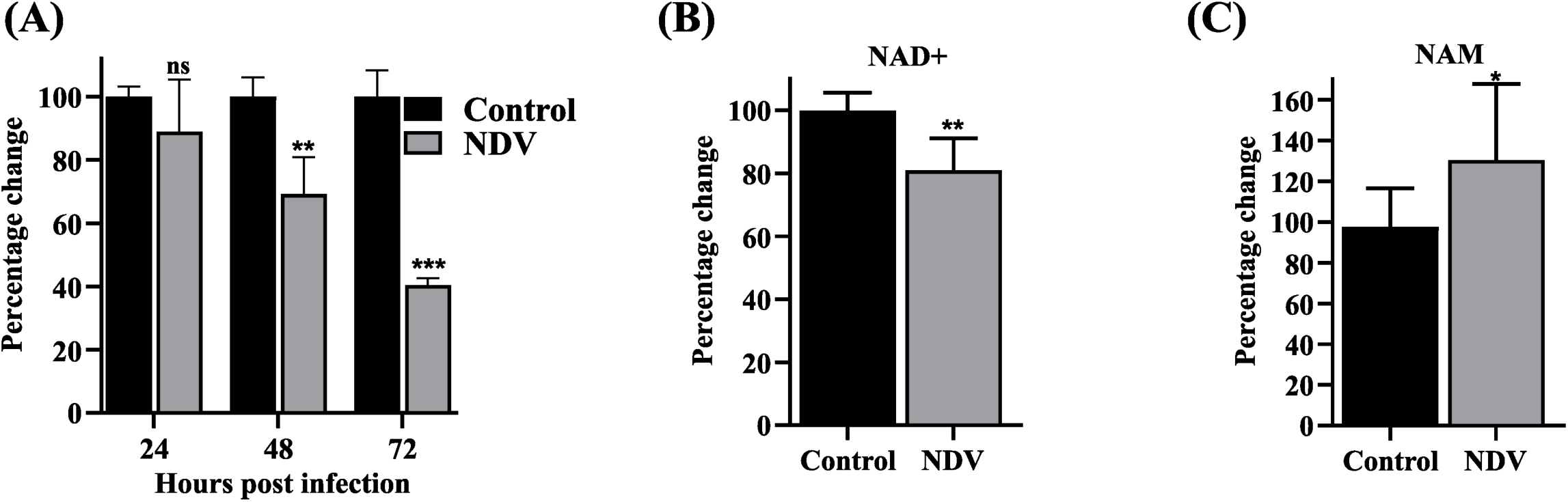
NAD^+^ and NAM estimation in NDV-infected cells and embryo tissues. Graph representing the concentration variation of NAD^+^ in control and NDV-infected DF-1 cells at different time points (A). Graph showing the variation in NAD^+^ concentration in NDV-infected embryo tissues (B). Graph showing the variation in NAM concentration in NDV-infected embryo tissues (C). Statistical analyses were performed using an unpaired two-tailed t-test, and the statistical significance among different groups was indicated as *, **, and *** where p < 0.05, p < 0.01, and p < 0.001, respectively.

## Discussion

Mitochondria are ubiquitous multifunctional organelles that assist the cells’ survival through metabolism and biosynthesis. Being a principal source of ROS in cells, mitochondria play a pivotal role in modulating oxidative stress and innate immune responses (66–68). Any abnormality in a typical cellular environment can cause an imbalance in ROS production-and-removal system, resulting in higher accumulation (69). ROS are often considered etiologic in inducing cellular injury by altering the cell’s redox state. A variety of viruses, including human immunodeficiency virus (HIV), dengue, influenza, and SARS-CoV-2, are reported to exploit or manipulate mitochondrial functions to induce oxidative stress, thus, promoting their replication (5, 6, 70–72). It brings attention to the crosstalk mechanism of oxidative stress and the pathogenesis of viral infections, which remains hither and thither. Oxidative stress typically triggers the host antiviral innate response; however, recent studies have described that elevated ROS levels may favor viral replication (70, 73–75).

Here, we tested the intrinsic impact of NDV infection on oxidative stress and intracellular ROS production. The results reflect the high generation of ROS temporally as the NDV infection cycle progressed. Simultaneously, comparing these elevated levels in infected cells with the positive inducer (DMF)-treated and ROS scavenger (PDTC and NAC)-treated cells confirm the presumption of ROS induction due to NDV infection. It was further validated via post-LiCl treatment, which brought down the ROS levels by inhibiting the NDV replication. To determine the plausible significance of the NDV-induced increased ROS levels in the pathogenesis, we studied the propensity of NDV to replicate in an oxidatively stressed environment. The initial results showed that DMF pre-incubated stressed cells had higher NDV RNA and protein expression, reflecting comparatively inflated virulence than in normal DF-1 cells. Considering the enhanced NDV replication, it was likely that the decrease in β-actin band intensity at 72 hrs in immunoblotting was attributable to NDV-induced cytopathic effect in addition to DMF treatment. The viral quantification assays also produced similar higher trends in virus titer. Additionally, PDTC attenuated the ROS generated in NDV-infected cells through direct scavenging, thus making a corresponding sharp decrease in NDV titer. In conjunction, these results confirm the involvement of intracellular ROS generation and indicate their importance in maintaining a stressed environment, thus helping the NDV replicate better.

Cellular response to the virus-imposed stress is mainly controlled by the universally conserved stress-responsive proteins, which are upregulated depending upon the type of stress the cell is experiencing (76). Viruses often manipulate the cellular response to maintain the oxidative stress environment throughout the infection cycle by inhibiting the synthesis and activity of antioxidant enzymes (63, 77–79). Therefore, we determined the expression patterns of crucial oxidative stress-responsive genes to examine the persistence of oxidative stress during infection. The suppression of Nrf2 and its downstream effector HO-1 suggested that NDV interferes with Nrf2/Keap1-signalling pathway, coming in concordance with the previous similar reports (80–82). Interestingly, SOD-1 activation was seen to be appreciable in the early stages of infection, insinuating that the infected cells might benefit from their activation to confer increased resistance against the virus-induced oxidative stress. However, the SOD-1 activity might be unfavorable for replication, and NDV therefore reduces its expression drastically, as shown previously (83). Taken together, the cellular gene expression profile data confirmed the modulation of oxidative stress-related genes post-NDV-infection, thus implying their implicit or explicit role in viral pathogenesis.

Additionally, expression profiles of sirtuin family genes were also studied, and expectedly, they are found to be differentially altered due to NDV infection. The magnitude of the variations in expression levels was strong enough to suggest their active participation in viral pathogenesis. In recent years, sirtuins have been mainly studied for their regulatory roles in cell metabolism, aging, and neurodegenerative diseases (36, 84–86). Moreover, numerous studies support their implication in mediating oxidative stress response to protect the cells from excessive ROS by directly deacetylating several key transcription factors, thus, regulating antioxidant gene expression (87–90). Hence, the prominent activation of SIRT1 and SIRT7 could be directly linked to the oxidative stress induced by NDV. Similar results are also reported showing their overexpression and constructive role in the progress of virus infection (45, 56, 91, 92). However, SIRT3 and SIRT5 were inhibited by NDV, which corroborates with the recent research work (93) showing the reprogramming of energy metabolism in infected cells after their degradation. The involvement of sirtuin proteins in stress-related pathways has unvaryingly strengthened the knowledge of viral pathogenesis, especially when the holes in our understanding of pathogenesis and host-response hindering strategies are perceived to be significant.

SIRT7 is an NAD+-dependent deacetylase located in the nucleus that regulates gene expression under oxidative stress conditions (53, 94, 95). In this study, we observed that SIRT7 protein is substantially upregulated in NDV-infected cells, complementing the transcriptional profile results. DMF and PDTC treatments showed the association of SIRT7 with oxidative stress; therefore, it is possible that expression enhancements shown here may be directed by NDV-induced stress to control its activity. Generally, the antiviral proteins are responsible for onsetting the antiviral response to perturb the infection. SIRT7 induced during virus infection may halt the progression of antiviral response, enabling the virus to replicate effectively. NDV infection in SIRT7-overexpressed cells manifested increased viral protein production than normal cells, thus confirming the hypothesis. These results collectively conveyed that SIRT7 expressed in stressed cells due to ROS accumulation plays a constructive role in NDV replication and the ROS scavenging consequentially decreased the virus yield. The subsequent experiments addressed the deacetylating activity of SIRT7 on cellular proteins.

Post-translational modifications regulate the cellular proteome by prompting structural changes in proteins, thus, affecting their functionality (96). Histone modifications, mainly acetylation and deacetylation, function as a specific transcription regulator governing cellular response (97). The addition and removal of acetyl functional groups to lysine residues can change the chromatin architecture and affect the chromatin’s open-and-closed state to regulate gene expression (98–100). Sirtuins, well-characterized class III histone deacetylases (HDACs) that depend on NAD^+^, can also influence multiple proteins other than histones (101, 102). In the present study, SIRT7-influenced levels of acetylated proteins were determined in NDV-infected cells. The temporal study reveals the variations in global acetylation profile due to NDV infection, which were further validated with SIRT7 overexpression showing complete abrogation. Correspondingly, a sharp drop in acetylated proteins was observed in cells when infected with different MOIs, with the most pronounced effect observed at higher concentrations of the virus. The heat-inactivation experiment suggests that the observed changes in acetylation levels were indeed affected by NDV replication in DF-1 cells. These observations of differential deacetylated proteins in infected cells provided information about NDV’s induced impediment of host protein production to adjourn the cellular defense mechanisms. The data we are reporting herein unveil the importance of SIRT7-driven deacetylation for viral pathogenesis.

Furthermore, detailed mechanistic studies were also performed with SIRT7 activity modulators. NAD-consuming enzymes, including sirtuin and poly-ADP-ribose polymerase (PARP) protein families, could metabolize NAD^+^ to produce 2’-O-Acetyl-ADP-ribose and NAM (103–105). NAD^+^ being a rate-limiting co-substrate for sirtuin enzymes, its restoration and scarcity could help control their deacetylation activity (84, 106). Predominantly NAD^+^ is synthesized through NMN (107, 108); thus, promoting NAD^+^ biosynthesis by NMN administration can effectively enhance SIRT7 activity (54), while NAM treatment can suppress the activity by compromising NAD^+^ levels via feedback inhibition (109). The β-NMN supplementation exhibited reduced acetylated proteins, which implied that NAD^+^ boosted concentrations induce SIRT7 activity. Moreover, we found that the NDV protein production was significantly increased when we treated the cells with β-NMN before infection. In contrast, NAM treatment negatively affects virulence, suggesting that NAD^+^ biosynthesis can control the SIRT7-mediated deacetylation of cellular proteins during the infection. These findings were further supported by the *in ovo* study, which aligns with those of cell culture results. Altogether, these observations suggest that the β-NMN supplementation not only restores normal NAD^+^ levels but also increases the virulence of NDV in the host. As could be seen from the results, NDV infection could overall lead to high SIRT7 activity, which suggests that NAD^+^ concentration may deplete post-infection due to an increased demand for deacetylating host proteins. The concentration of NAD+ was determined using an HPLC system to evaluate this possibility. The depletion of NAD^+^ levels with time in infected cells clearly supports the presumptions made and emphasizes the importance of NAD^+^ decrease as a common trigger of virus-associated cytopathic effects. Consistent with these findings, embryo tissues also provide compelling support for NAD^+^ scarcity while showing boosted levels of NAM, which might be the consequence of SIRT7-driven NAD^+^ metabolism to deacetylate the proteins and generate NAM as a by-product.

Taken together, the data obtained in the study suggest that high intracellular ROS generation and the resulting oxidative stressed environment are necessary for effective NDV replication. Moreover, the NDV-modulated stress elicits several transcriptional and translational outputs that activate stress-responsive proteins, which the virus utilizes for better replication. NDV-induced oxidative stress drives the production of SIRT7 in DF-1 cells, leading to the deacetylation of host proteins. The utilization of SIRT7-mediated deacetylation could be a crucial underlying strategy employed by NDV to thrive in its host. The notable result of our study confirms that SIRT7 once activated in DF-1 cells due to mitochondrial damage and ROS accumulation, supports virus infection. Lastly, evaluating the drugs and molecules that can modulate sirtuin activity will be essential in developing an effective antiviral strategy to control the disease.

## Acknowledgments

We thank Ms. Sandhya Sekhar and Mr. Manish Ghumnani (Bioprocess Analytical Technology laboratory) for their help in analyzing the HPLC data. The virus research in our laboratory is currently supported by the Department of Biotechnology, India (BT/PR41246/NER/95/1685/2020) and the Department of Health Research, Government of India (NER/71/2020-ECD-I).

---

**Figure S1.**
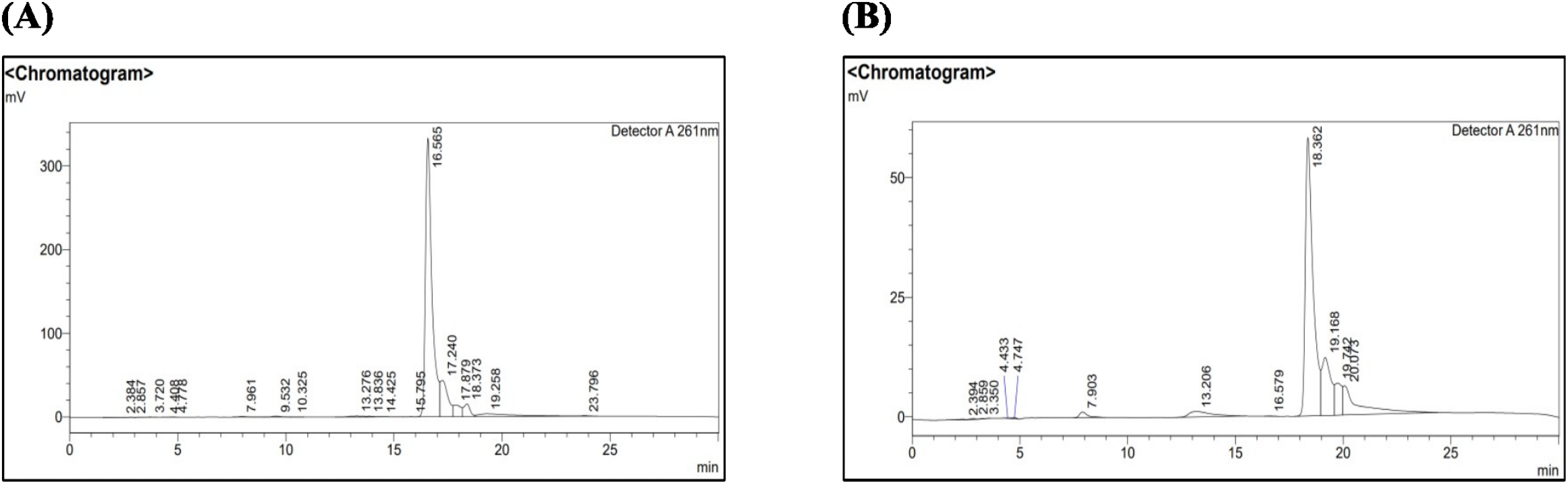
HPLC chromatograms obtained for NAD^+^ and NAM standard solutions (100 μM) (A)(B).

**Figure S2.**
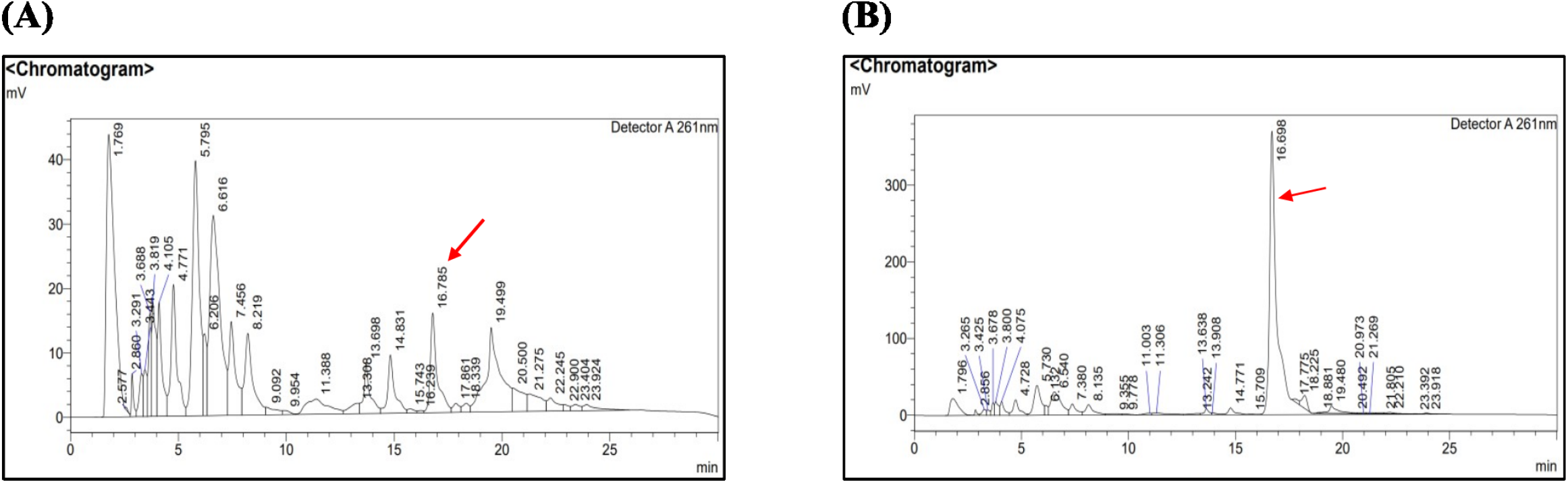
HPLC chromatograms obtained for DF-1 cells with and without NAD^+^ supplementation (A)(B).

**Figure S3.**
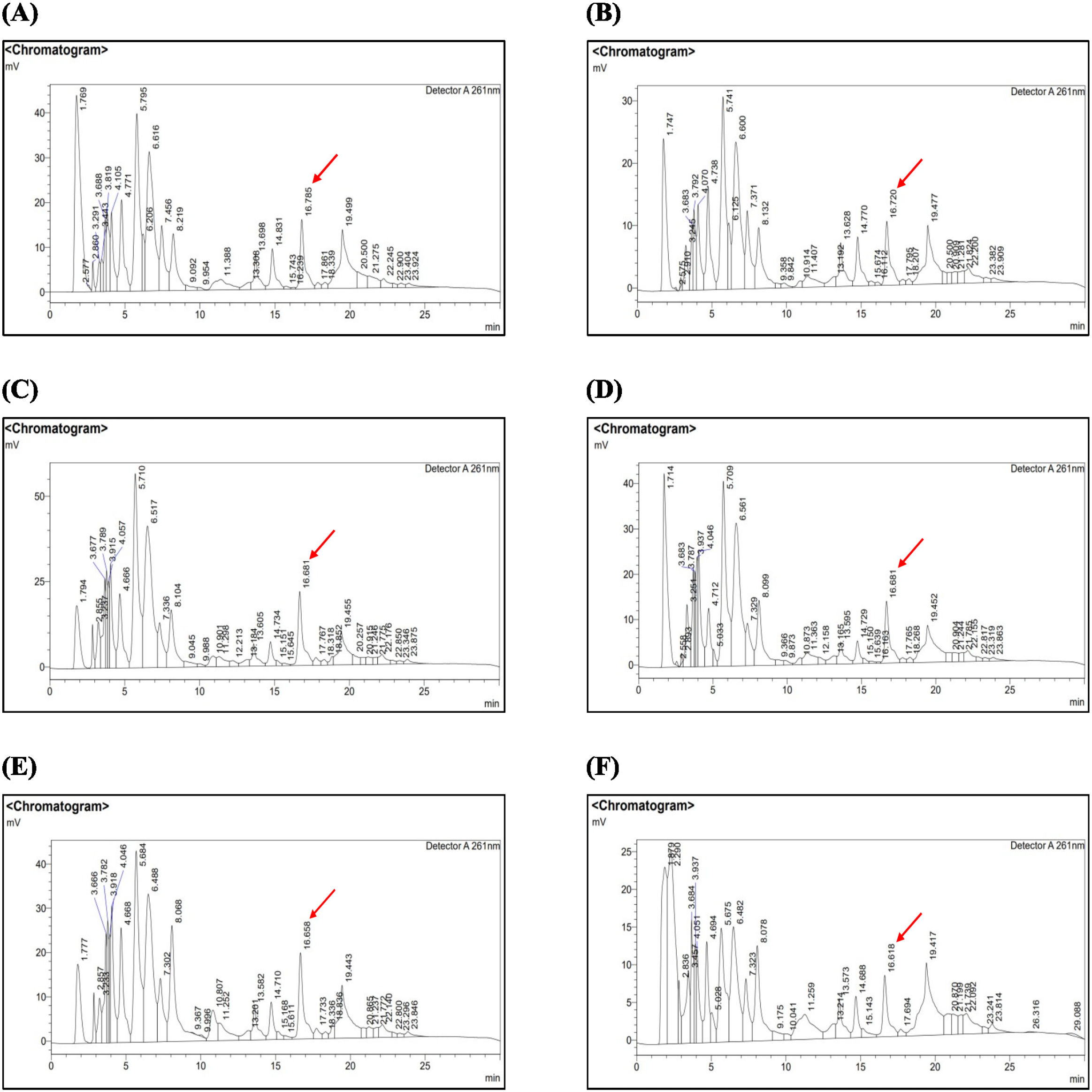
HPLC chromatograms obtained for control and NDV-infected DF-1 cell samples at 24 (A)(B), 48 (C)(D), and 72 (E)(F) hours, highlighting the NAD^+^ peaks.

